# InterClone: Store, Search and Cluster Adaptive Immune Receptor Repertoires

**DOI:** 10.1101/2022.07.31.501809

**Authors:** Jan Wilamowski, Zichang Xu, Hendra S Ismanto, Songling Li, Shunsuke Teraguchi, Mara Anais Llamas- Covarrubias, Xiuyuan Lu, Sho Yamasaki, Daron M Standley

## Abstract

B and T cell receptor repertoire data has the potential to fundamentally change the way we diagnose and treat a wide range of diseases. However, there are few resources for storing or analyzing repertoire data. InterClone provides tools for storing, searching, and clustering repertoire datasets. Efficiency is achieved by encoding the complementarity-determining regions of sequences as mmseqs2 databases. Single chain search or cluster results can be merged into paired (alpha-beta or heavy-light) results for analysis of single-cell sequencing data. We illustrate the use of InterClone with two recently reported examples: 1) searching for SARS-CoV-2 infection-enhancing antibodies in bulk COVID-19 and healthy donor repertoires; 2) identification of SARS-CoV-2 specific TCRs by clustering paired and bulk sequences from COVID-19, BNT162b2 vaccinated and healthy unvaccinated donors. The core functions of InterClone have been implemented as a web server and integrated database (https://sysimm.org/interclone). All source code is available upon request.

## Introduction

Adaptive immune responses are mediated by B and T lymphocytes, which harbor rearranged antigen-receptor gene segments. The set of B cell receptors (BCRs) or T cell receptors (TCRs) in a given donor at a given time is known as the donor’s *repertoire*. Because each repertoire encodes a record of past and present antigen exposures, which in turn influences responses to future antigen encounters, a great amount of interest has been paid to developing methods that can associate BCRs or TCRs with their targeted antigens or with a donor’s health/disease status. To facilitate these efforts, Adaptive Immune Receptor Repertoire (AIRR) standards for describing repertoire data have been established (Rubelt, et al., 2017).

## Results

### Storing data

AIRR-formatted datasets can be uploaded to InterClone by anyone, without the need for an account. Data visibility is controlled by the user: private data is visible only to the registered and logged in owner of an account, while public data is visible to everyone.

### Searching

A query sequence can be searched against a group of stored datasets at the amino acid sequence level, using user-defined similarity thresholds for each complementarity-determining region (CDR). To illustrate the search function, we utilized 11 recently-reported SARS-CoV2 infection enhancing antibodies (Li, et al., 2021; Liu, et al., 2021). These antibodies have apparently emerged from different germline genes and possess highly diverse CDRH3 amino acid sequences but target an overlapping patch of residues on the N-terminal domain (NTD) of the SARS-CoV2 spike protein (Table S1). In order to assess the frequency of these 11 antibody families in the general population, we searched 4.5 million unique BCR heavy chain sequences acquired from COVID-19 donors, as well as 5.1 million unique BCR heavy chain sequences from three healthy unvaccinated donors (Table S2). Details of the search are provided in Methods. As shown in Figure 1A, InterClone found a total of 374 hits in the COVID-19 dataset for 8 of the 11 enhancing antibodies. This number is somewhat higher than the 232 hits for 7 of the enhancing antibodies found in the healthy datasets. Moreover, after considering identical clones, the number of hits in the COVID-19 donors (7259) was significantly higher than in the healthy unvaccinated donors set (1623), suggesting clonal expansion in infected donors. As a reference, we also ran CompAIRR (Rognes, et al., 2022), which uses V and J gene matching along with the CDRH3 edit distance to identify similar sequences, on the same inputs. While both tools could find hits, InterClone searched much more broadly. This is important because we have observed that the sequence identities with a family of true enhancing antibodies can be as low as 80% in CDRH1-2 and 70% in CDRH3 (Ismanto, et al., 2022). In such a diverse search space we found that InterClone found more relevant results at comparable thresholds than CompAIRR (Table S3). Details of the CompAIRR search procedure are given in Methods.

**Fig. 1.**
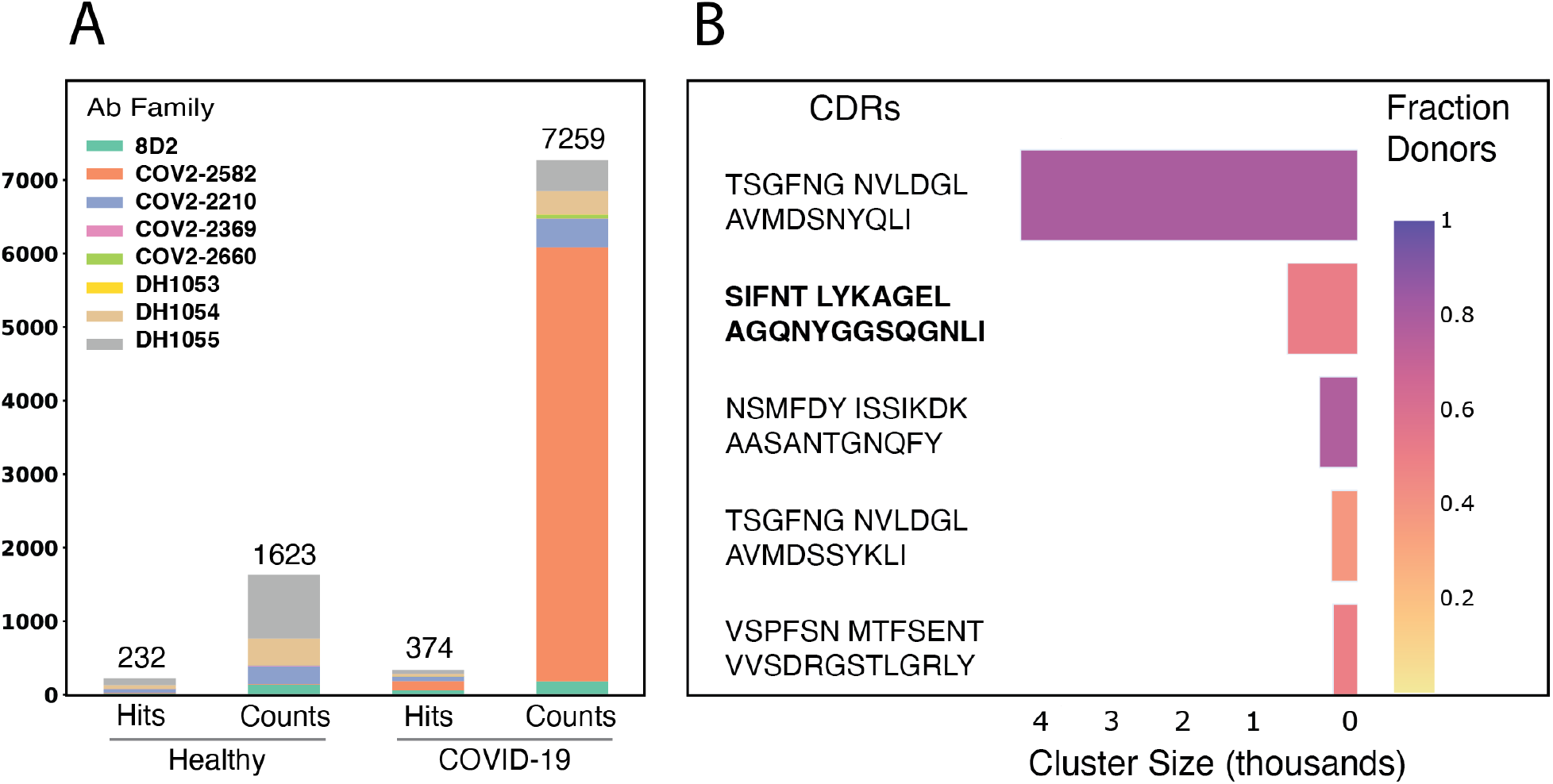
Searching and clustering. A, Search hits and clone counts for 8 SARS-CoV-2 infection-enhancing antibodies. B, Cluster results for 98 COVID-19 donor TCRs, with darker colors indicating higher cluster occupancy by donors.

### Clustering

To cluster a group of datasets by overall CDR sequence similarity, InterClone uses mmseqs cluster, which scales approximately linearly with input data size and thus can readily handle multiple repertoires efficiently. Details of the Clustering are provided in Methods. We clustered TCR alpha chain data collected from 98 COVID-19 donors using a 90% sequence identity threshold. Figure 1B shows the resulting clusters, ranked by size. The majority of large public clusters corresponded to Mucosal-associated invariant T (MAIT)-like sequences, which recognize metabolites. However, the second-largest cluster contained a CDRA3 motif that matched that reported previously to recognize S protein-derived peptides (Mudd, et al., 2022). Similar results were observed in a second TCRA dataset corresponding to single T cells acquired from BNT162b2 vaccinated donors and cultured in S protein-derived peptides (Figure S2A), but not in PBMCs acquired pre-vaccination (Figure S2B) or healthy unvaccinated donors (Figure S1B). We also clustered TCRA data with CompAIRR using various thresholds and observed consistent results: a single cluster containing the S-protein specific motif as well as many invariant alpha chain motifs (Figure S3A).

### Merging

The search and cluster functions both operate on the single chain level. Paired (BCR light-heavy, TCR alpha-beta) results can be prepared using the merge feature for single cell sequence data, as detailed in Methods. To illustrate merging results from individual chains, we clustered the TCR beta chains from the BNT162b2 vaccinated donor data, then represented their linkage for the S-protein specific cluster (Figure S4). We found that the beta chain sequences were much more diverse, as reported previously (Mudd, et al., 2022), but that patterns in CDRB3 could nevertheless be readily observed.

## Methods

InterClone provides flexible methods for storing, searching, and clustering repertoire data, along with a central repository of repertoire sequences. To store a group of repertoires, the inputs must consist of one or more AIRR-formatted files in a zip archive. Procedures for preparing AIRR-formatted files from raw data are described in Suppl Text T1. To enable efficient downstream analyses, the amino acid sequences are first decomposed into the three CDRs, using ANARCI (Dunbar and Deane, 2016) with IMGT numbering. The CDRs are then concatenated into a single pseudosequence and encoded as mmseqs2 databases (Steinegger and Soding, 2017). The backend command-line interface is highly customizable and described in more detail in the source repository.

### InterClone search procedure

Given a set of three CDR similarity thresholds, the lowest threshold (min_sid) is used in the initial mmseqs search, along with the following parameters:

- -s 2 (sensitivity)
- --mask 0 (no low complexity masking)
- -a (add backtrace string, for convertalis)
- --min-seq-id min_sid (minimum sequence identity)
- -c cov (minimum residue coverage fraction)

The results are then filtered by the user-defined CDR similarity and coverage thresholds on the fly. For the search for enhancing antibodies, we used the following thresholds:

- CDR1: 80%
- CDR2: 80%
- CDR3: 70%
- Coverage: 80%

### CompAIRR search procedure

For comparison, we ran CompAIRR v1.7.0 on the same inputs with the following parameters:

- --existence (equivalent search function)
- --ignore-counts (do not use the duplicate_count field)
- --differences d (maximum number of allowed mutations)
- --pairs (output matching pairs)

While our chosen Interclone CDR3 sequence similarity threshold of 0.7 (i.e. 70%) is applied to each sequence separately, a fixed value has to be selected for CompAIRR. To determine the value of d that gives the closest result to Interclone, we calculated the average length of CDR3 sequences in our dataset. For the COVID-19 data, this is 14.9 (median 15) and for the healthy data, this is 15.4 (median 15). The maximum allowed deviation from this is 1-0.7 = 30%, which gives the maximum edit distance of 4.5-4.6. Thus, we ran CompAIRR with the values 4 and 5 for the parameter d. It is noteworthy that due to their different threshold definitions (relative vs absolute), the two tools will never produce the exact same results. Furthermore, Interclone allows flexible cutoff values for the other two CDRs whereas CompAIRR will filter by requiring exact matches of the J and V gene names. This fundamentally compares different parts of the full-length sequence, again yielding distinct results (see Table S3). Due to the different search criteria, a large proportion of hits for a specific parameter set are unique to either method. We would like to point out that simply returning more hits is not necessarily useful, as the result count can always be increased by lowering thresholds/increasing allowed differences. Each tool can produce valid results depending on the use case, that is, depending on whether CDR sequence or V/J gene name comparison is more appropriate. However, one limitation of CompAIRR is that indels (i.e. insertions or deletions) can only be accounted for when using d=1. Due to the large variability in CDR3 length, this limitation prevents CompAIRR from recognizing similar sequences. As discussed above, values of d much higher than 1 are required for diverse datasets.

### InterClone clustering procedure

The input databases are first combined into one database. Next, mmseqs cluster is invoked with the following parameters:

- --min-seq-id min_sid (minimum sequence identity)
- -c cov (minimum residue coverage fraction)

From the results, a TSV file is generated using mmseqs createtsv. Since these results only contain sequence identifiers without metadata, they are merged with the original input files. This allows for the downstream analysis of any metadata contained in the source data, e.g. donor assignment, time of sample extraction, etc.

### Procedure for merging paired single-chain results

The pseudo sequence preparation, search and clustering functions handle paired input data transparently by processing them separately and storing the outputs in their own folders. If the user is interested only in those results that appear in both heavy and light (or alpha and beta) subsets, the results can be merged. An identifier is required to find pairs, by default this is the clone_id column in each AIRR file. The results are first filtered to ensure the existence of both heavy and light (or alpha and beta) chains. However, further duplication (e.g. two light chains) is not removed so it is important to choose an appropriate identifier column. This identifier is not required to be globally unique; instead, it should only be unique per AIRR file.

## Acknowledgements

The authors would like to thank beta users for their valuable feedback.

