## Supplementary Materials for "InterClone: Store, Search and Cluster Adaptive Immune Receptor Repertoires"

### Supplementary Data

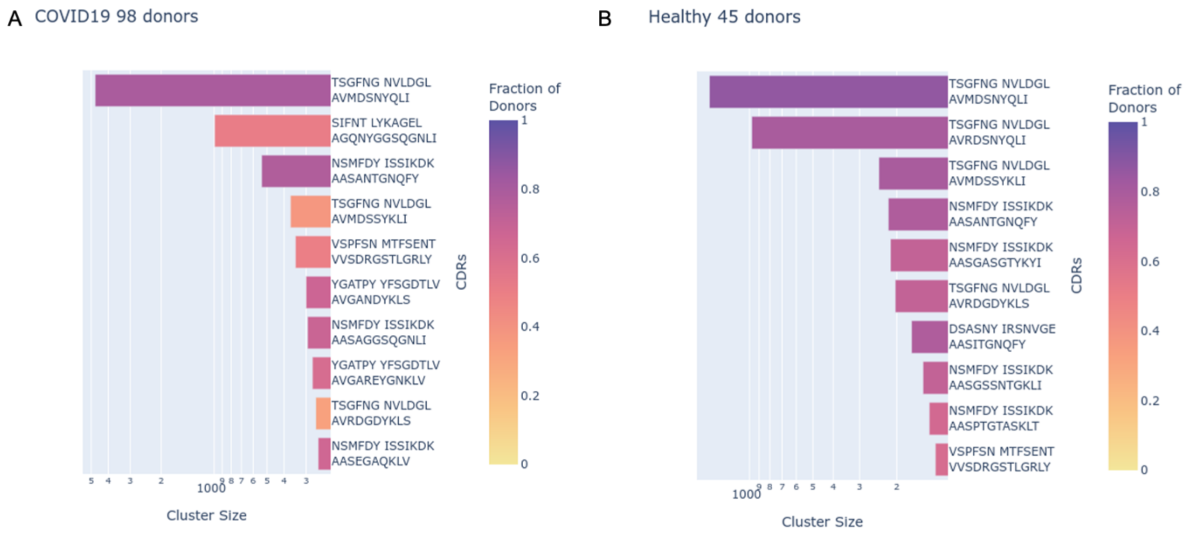


**Figure S1. InterClone TCR alpha chain Cluster results for COVID-19 (A) and Healthy (B) donors.** Cluster sizes are shown on a log scale. CDR sequences for cluster representatives are shown to the right of the bars. The color palette, which indicates donor coverage, is the same as in Figure 1B. The Public TCRA motif in COVID data is given by SIFNT/LYKAGEL/A[G/A/V]XNYGGSQGNLI

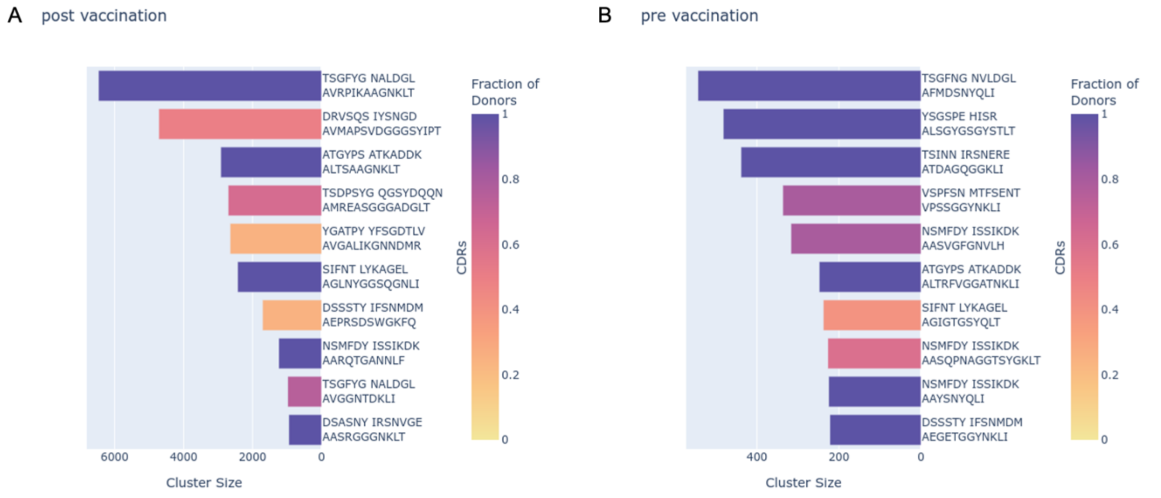


**Figure S2. InterClone TCR alpha chain Cluster results for post-vaccinated (A) and pre-vaccinated (B) donors.** For vaccinated donors, PBMCs were cultured with S protein peptides. Format is the same as Figure S1.

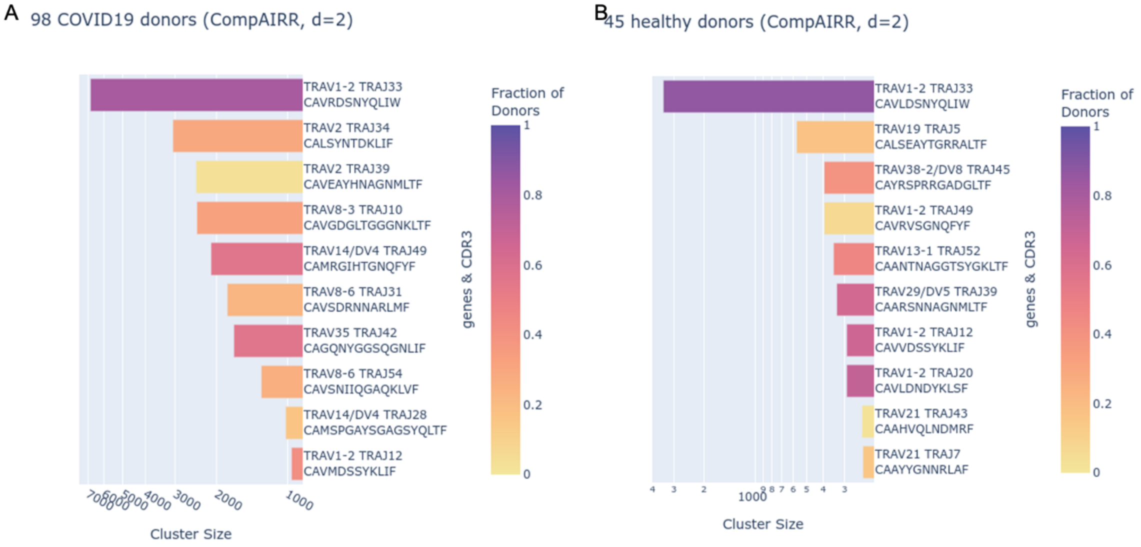


**Figure S3. CompAIRR TCR alpha chain Cluster results for 98 COVID-19 (A) and (B) 45 healthy donors.** CompAIRR was run with d=2. The public TCRA motif in the COVID data is given by TRAV35 / TRAJ42 / CA[G/A/V]XNYGGSQGNLIF. The data source and figure format are the same as that of Figure S1.

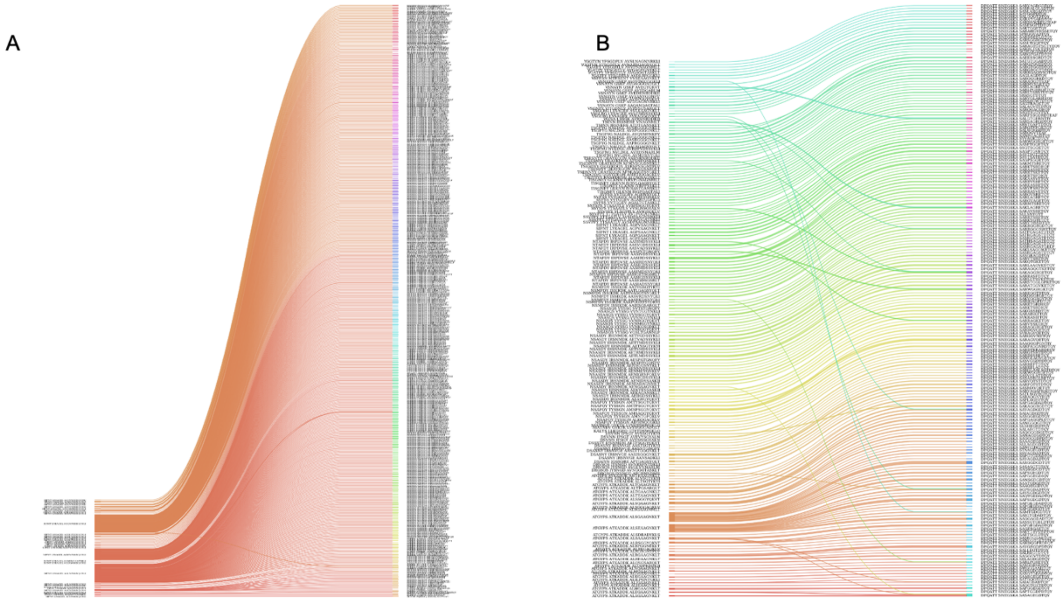


**Figure S4. Example of merging TCR alpha and beta chain cluster results.** A, The CDRs of the largest alpha cluster (left side) are shown with their paired beta chain CDRs (right side). B, The CDRs of the largest beta cluster (right side) are shown with their paired alpha chain CDRs (left side). The data source is the same as that for Figure S2.

#### Table S1. Known enhancing antibody sequence and gene usage

| Name | $V_H$ gene | $J_H$ gene | CDRH3 aa | $V_L$ gene | $J_L$ gene | CDRL3 aa |
|---|---|---|---|---|---|---|
| 8D2 | IGHV3-7 | IGHJ3 | ARDWDYDILTGSWFGAFDI | IGKV1-17 | IGKJ4 | LQHNSYPLT |
| COV2-2210 | IGHV3-30-3 | IGHJ4 | ARDQEWFRELFLFDY | IGKV1-12 | IGKJ3 | QQANSFPPT |
| COV2-2369 | IGHV3-30 | IGHJ4 | AKDFGGDNTAMVEYFFDF | IGKV1-5 | IGKJ1 | QQYNSYSPT |
| COV2-2490 | IGHV3-7 | IGHJ4 | ARDPYDLYGDYGGTFDY | IGKV1-5 | IGKJ4 | QQYNSYSLT |
| COV2-2582 | IGHV7-4-1 | IGHJ6 | ARDQDSGYPTYYYYYMDV | IGKV2D-29 | IGKJ4 | MQSIQPPLT |
| COV2-2660 | IGHV3-13 | IGHJ6 | ARADPYQLLGQHYYYGMDV | IGKV3-20 | IGKJ5 | QQYGSSPLIT |
| DH1052 | IGHV1-69-2 | IGHJ4 | ATSSGPSRLCGGGSCYHSFDY | IGKV3-20 | IGKJ1 | QQYGSSPTWT |
| DH1053 | IGHV3-43 | IGHJ4 | AKAKDPYTEYFDY | IGKV1-5 | IGKJ4 | QQYYIYSLS |
| DH1054 | IGHV3-53 | IGHJ3 | ARGDIVGATWDPAFDI | IGLV3-21 | IGLJ2 | QVWDTSSDHSVV |
| DH1055 | IGHV2-70 | IGHJ4 | ARINAYSSSWPTFDY | IGKV3-20 | IGKJ1 | QQYGSSSWT |
| DH1056 | IGHV4-39 | IGHJ5 | ARSSSGFSYDTPLDP | IGKV1-17 | IGKJ5 | LQHNSYPIT |

#### Table S2: InterClone datasets used for BCR search and TCR clustering examples

| Input | Dataset | Size | References |
|---|---|---|---|
| Enhancing antibody queries BCR | enhancing-antibodies | 11 | (Li et al. 2021; Liu et al. 2021) |
| COVID-19 target BCR | Kim-2021 | 4984980 | (Kim et al. 2021) |
| Healthy Targets BCR | Ghraichy-2020<br>Gidoni-2019<br>Meng-2017 | 1932766<br>1687626<br>1493033 | (Ghraichy et al. 2020; Gidoni et al. 2019; Meng et al. 2017) |
| TCR vaccinated | TCR-postvac | 65758 | (Lu 2022) |
| TCR pre-vaccinated | TCR-prevac | 6775 | (Lu 2022) |
| TCR COVID19 | Zhang-2020-a<br>Zhang-2020-b<br>Wen-2020<br>Sureshchandra-2021<br>Ramaswamy-2021<br>Notarbartolo-2021<br>Meckiff-2020<br>Liao-2020<br>Bieberich-2021<br>Bacher-2020 | 13360<br>22436<br>10287<br>9203<br>41088<br>98471<br>86083<br>2749<br>82472<br>22036 | (Notarbartolo et al. 2021; Ramaswamy et al. 2021; Sureshchandra et al. 2021; Wen et al. 2020; Zhang, Gan, et al. 2020; Zhang, Wang, et al. 2020) |

| | | | |
|---|---|---|---|
| TCR Healthy | Zhang-2020-a<br>Zhang-2020-b<br>Wen-2020<br>Sureshchandra-2021<br>Ramaswamy-2021<br>Notarbartolo-2021<br>Bacher-2020<br>Luo-2022<br>Gao-2022 | 11294<br>4043<br>6185<br>8681<br>47563<br>10285<br>1204<br>17461<br>55919 | (Bacher et al. 2020; Gao et al. 2022; Luo et al. 2022; Notarbartolo et al. 2021; Ramaswamy et al. 2021; Sureshchandra et al. 2021; Wen et al. 2020; Zhang, Gan, et al. 2020; Zhang, Wang, et al. 2020) |

**Table S3: Number of hits for two search methods, showing overlap and result set differences**

| Dataset | Max. differences (d) | Shared hits | Only Interclone | Only CompAIRR |
|---|---|---|---|---|
| COVID-19 | 4 | 19 | 315 | 3 |
| | 5 | 24 | 310 | 42 |
| Healthy | 4 | 6 | 218 | 1 |
| | 5 | 6 | 218 | 27 |

**Text T1: Preparing AIRR-formatted files**

InterClone requires AIRR-formatted files, which are tab-delimited files with a number of specific column headers. In order to use InterClone, the following headers are required:

- sequence_id, containing a unique identifier for a sequence entry
- sequence_aa, containing the complete amino acid sequence
- v_call, containing the assigned V gene name for chain filtering
- clone_id, containing a unique identifier for each clone, used for paired result merging

Input files can be prepared from raw FASTA, 10X CellRanger, Illumina MIRA and MiXCR-formatted data. An import script is provided for each of these data sources. They can be found in the src/dataimport/ folder in the source code repository.

In the case of FASTA-formatted data, the user provides one or more input files containing full length amino acid sequences, along with the chain type of these sequences.

In the case of raw MiXCR TSV outputs, AIRR files are prepared as follows:

1. Conjugate the full-length amino acid sequence by merging the CDR and framework regions from 'aaSeqImputedFR1', 'aaSeqImputedCDR1', 'aaSeqImputedFR2', 'aaSeqImputedCDR2', 'aaSeqImputedFR3', 'aaSeqImputedCDR3' and 'aaSeqImputedFR4'. Discard sequences containing gaps or stop codons in these regions.
2. The columns 'cloneId', 'allVHitsWithScore', 'allJHitsWithScore', 'aaSeqImputedCDR3', 'allCHitsWithScore’, and ‘cloneCount' from the MiXCR output file are renamed to 'clone_id', 'v_call', 'j_call', 'cdr3', 'c_call' and 'clone_count', respectively.

In the case of 10X CellRanger output, three files are required: 'airr_rearrangement.tsv', 'clonotypes.csv' and 'all_contig_annotations.csv'. To prepare AIRR formatted files, the following steps are taken:

1. Examine the quality of each clone: Only clones that contain one paired heavy/light or alpha/beta chain are used. Clones containing multiple chains or single chains are discarded. The frequency of each clone taken from ‘clonotypes.csv’ and saved as 'clone_count' in the AIRR file.

2. Select one of the contigs that share the same clonotype and use the columns 'cell_id', 'v_call', 'j_call', 'cdr3', 'c_call' and amino acid sequence for the AIRR file.

InterClone filters inputs by chain type using the ‘v_call’ column. If both heavy and light chain data is included in an input file, the pseudo-sequences of each chain will be stored separately and processed individually in the next steps.
Most software tools for repertoire analysis have ways of handling duplicated sequences. Usually, these would be grouped under a common clonotype with a unique identifier. Since the proper processing of these duplicates depends greatly on the metadata, InterClone does not try to automatically deduce these relationships. For example, duplicated sequences within the same donor should be eliminated and recorded via a column like clone_count. On the other hand, duplicated sequences among multiple donors could be considered a signal of interest in cluster analysis.

Liu, Yafei, Noriko Arase, Jun-ichi Kishikawa, Mika Hirose, Songling Li, Asa Tada, Sumiko Matsuoka, Akemi Arakawa, Kanako Akamatsu, Chikako Ono, Hui Jin, Kazuki Kishida, Wataru Nakai, Masako Kohyama, Atsushi Nakagawa, Junichi Takagi, Yoshiaki Yamagishi, Hironori Nakagami, Atsushi Kumanogoh, Yoshiharau Matsuura, Daron M. Standley, Takayuki Kato, Masato Okada, Manabu Fujimoto, and Hisashi Arase. 2021. 'Acquisition of resistance to wild-type spike-immune sera by emerging SARS-CoV-2 variants', *Research Square*.

Lu, X. 2022. 'in prep'.
Luo, Oscar Junhong, Wen Lei, Guodong Zhu, Zhiyao Ren, Yudai Xu, Chanchan Xiao, Hongyi Zhang, Junxiang Cai, Zhiping Luo, Lijuan Gao, Jun Su, Lei Tang, Wei Guo, Huanxing Su, Zhang-Jin Zhang, Evandro Fei Fang, Yijun Ruan, Sean Xiao Leng, Zhenyu Ju, Huiling Lou, Junling Gao, Nan Peng, Jie Chen, Zhijun Bao, Feng Liu, and Guobing Chen. 2022. 'Multidimensional single-cell analysis of human peripheral blood reveals characteristic features of the immune system landscape in aging and frailty', *Nature Aging*, 2: 348-64.
Meng, W., B. Zhang, G. W. Schwartz, A. M. Rosenfeld, D. Ren, J. J. C. Thome, D. J. Carpenter, N. Matsuoka, H. Lerner, A. L. Friedman, T. Granot, D. L. Farber, M. J. Shlomchik, U. Hershberg, and E. T. Luning Prak. 2017. 'An atlas of B-cell clonal distribution in the human body', *Nat Biotechnol*, 35: 879-84.
Notarbartolo, S., V. Ranzani, A. Bandera, P. Gruarin, V. Bevilacqua, A. R. Putignano, A. Gobbini, E. Galeota, C. Manara, M. Bombaci, E. Pesce, E. Zagato, A. Favalli, M. L. Sarnicola, S. Curti, M. Crosti, M. Martinovic, T. Fabbris, F. Marini, L. Donnici, M. Lorenzo, M. Mancino, R. Ungaro, A. Lombardi, D. Mangioni, A. Muscatello, S. Aliberti, F. Blasi, T. De Feo, D. Prati, L. Manganaro, F. Granucci, A. Lanzavecchia, R. De Francesco, A. Gori, R. Grifantini, and S. Abrignani. 2021. 'Integrated longitudinal immunophenotypic, transcriptional and repertoire analyses delineate immune responses in COVID-19 patients', *Sci Immunol*, 6.
Ramaswamy, A., N. N. Brodsky, T. S. Sumida, M. Comi, H. Asashima, K. B. Hoehn, N. Li, Y. Liu, A. Shah, N. G. Ravindra, J. Bishai, A. Khan, W. Lau, B. Sellers, N. Bansal, P. Guerrerio, A. Unterman, V. Habet, A. J. Rice, J. Catanzaro, H. Chandnani, M. Lopez, N. Kaminski, C. S. Dela Cruz, J. S. Tsang, Z. Wang, X. Yan, S. H. Kleinstein, D. van Dijk, R. W. Pierce, D. A. Hafler, and C. L. Lucas. 2021. 'Immune dysregulation and autoreactivity correlate with disease severity in SARS-CoV-2-associated multisystem inflammatory syndrome in children', *Immunity*, 54: 1083-95 e7.
Sureshchandra, S., S. A. Lewis, B. M. Doratt, A. Jankeel, I. Coimbra Ibraim, and I. Messaoudi. 2021. 'Single-cell profiling of T and B cell repertoires following SARS-CoV-2 mRNA vaccine', *JCI Insight*, 6.
Wen, Wen, Wenru Su, Hao Tang, Wenqing Le, Xiaopeng Zhang, Yingfeng Zheng, Xiuxing Liu, Lihui Xie, Jianmin Li, Jinguo Ye, Liwei Dong, Xiuliang Cui, Yushan Miao, Depeng Wang, Jiantao Dong, Chuanle Xiao, Wei Chen, and Hongyang Wang. 2020. 'Immune cell profiling of COVID-19 patients in the recovery stageby single-cell sequencing', *Cell Discovery*, 6: 31.
Zhang, Fan, Rui Gan, Ziqi Zhen, Xiaoli Hu, Xiang Li, Fengxia Zhou, Ying Liu, Chuangeng Chen, Shuangyu Xie, Bailing Zhang, Xiaoke Wu, and Zhiwei Huang. 2020. 'Adaptive immune responses to SARS-CoV-2 infection in severe versus mild individuals', *Signal Transduction and Targeted Therapy*, 5: 156.
Zhang, J. Y., X. M. Wang, X. Xing, Z. Xu, C. Zhang, J. W. Song, X. Fan, P. Xia, J. L. Fu, S. Y. Wang, R. N. Xu, X. P. Dai, L. Shi, L. Huang, T. J. Jiang, M. Shi, Y. Zhang, A. Zumla, M. Maeurer, F. Bai, and F. S. Wang. 2020. 'Single-cell landscape of immunological responses in patients with COVID-19', *Nat Immunol*, 21: 1107-18.
